# Overlapping roles of JIP3 and JIP4 in promoting axonal transport of lysosomes in human iPSC-derived neurons

**DOI:** 10.1101/2020.06.13.149443

**Authors:** Swetha Gowrishankar, Lila Lyons, Nisha Mohd Rafiq, Agnes Roczniak-Ferguson, Pietro De Camilli, Shawn M. Ferguson

## Abstract

The dependence of neurons on microtubule-based motors for the movement of lysosomes over long distances raises questions about adaptations that allow neurons to meet these demands. Recently, JIP3/MAPK8IP3, a neuronally enriched putative adaptor between lysosomes and motors, was identified as a critical regulator of axonal lysosome abundance. In this study, we establish a human induced pluripotent stem cell (iPSC)-derived neuron model for the investigation of axonal lysosome transport and maturation and show that loss of JIP3 results in the accumulation of axonal lysosomes and the Alzheimer’s disease-related amyloid precursor protein (APP)-derived Aβ42 peptide. We furthermore reveal an overlapping role of the homologous JIP4 gene in lysosome axonal transport. These results establish a cellular model for investigating the relationship between lysosome axonal transport and amyloidogenic APP processing and more broadly demonstrate the utility of human iPSC-derived neurons for the investigation of neuronal cell biology and pathology.

## Introduction

The endo-lysosomal system plays key house-keeping roles in all cells, but neurons are particularly dependent on efficient lysosome function owing to their extreme size, polarity and post-mitotic state (Ferguson, 2019). Indeed, rare loss-of-function mutations in multiple genes encoding lysosome proteins result in lysosome storage diseases, which frequently manifest with severe neurological and neurodegenerative pathologies (Sun, 2018; Marques and Saftig, 2019; Ballabio and Bonifacino, 2020). More recently, advances in understanding the genetics of neurodegenerative diseases associated with aging such as Alzheimer’s disease, Parkinson’s disease, amyotrophic lateral sclerosis and frontotemporal dementia, have identified a number of genes encoding endo-lysosomal proteins as modulators of disease risk (Lambert *et al*., 2013; Guerreiro and Hardy, 2014; Chang *et al*., 2017; Klein and Mazzulli, 2018; Wallings *et al*., 2019). Human genetics thus points to the importance of optimal lysosome function for the maintenance of neuron health and raises fundamental questions about how the endo-lysosomal system is adapted to the unique demands of neurons.

Primary cultures of embryonic or neonatal rodent neurons are a well-established model for the investigation of neuronal cell biology (Barnes and Polleux, 2009; Banker, 2018). In particular, primary cultures of cortical and hippocampal neurons from transgenic and knockout mice have been widely used to investigate the functions of specific genes in neurons. Although this has been a tremendously valuable model system, it has several significant limitations. These include practical issues such as the time required for establishing mouse colonies for each new mutation and the ongoing mouse breeding and genotyping required to provide embryos/pups for each new primary culture. Furthermore, combining multiple mutations into one mouse at minimum requires lengthy breeding schemes and often has negative impacts on mouse viability. Excitingly, the development of human induced pluripotent stem cells (iPSCs), combined with highly efficient protocols for their differentiation into defined neuronal cell types along with CRISPR-Cas9-dependent genome editing tools has opened up new opportunities for investigating the impact of targeted genetic perturbations on neuronal cell biology (Dolmetsch and Geschwind, 2011; Zhang *et al*., 2013; Paquet *et al*., 2016; Wang *et al*., 2017; Fernandopulle *et al*., 2018; Tian *et al*., 2019). In particular, the development of robust neuronal differentiation protocols based on transcription factor over-expression now allows for the generation of human neuronal cultures with unprecedented speed and consistency (Zhang *et al*., 2013; Fernandopulle *et al*., 2018).

In this study, we used human iPSC-derived neurons to investigate the relationships between lysosome axonal transport and Alzheimer’s disease pathology. This research direction builds on longstanding reports of lysosome-filled axonal dilations around Aβ deposits at amyloid plaques (Terry *et al*., 1964; Cataldo and Nixon, 1990; Nixon *et al*., 2005; Condello *et al*., 2011; Dikranian *et al*., 2012; Gowrishankar *et al*., 2015). Although this relationship between axonal lysosome accumulations and amyloid plaques has been known for many years, it has been challenging to define the relevance of these lysosomes to the broader disease pathology due to the lack of tools to selectively alter the abundance of axonal lysosomes. Recent progress on this topic came from observations that axonal lysosomes accumulated in neurons from JIP3 mutant mice and that JIP3 haploinsufficiency had a major impact on amyloid plaques (Gowrishankar *et al*., 2017). JIP3 is thought to act as an adaptor between lysosomes and the dynein motor; a function supported by axonal accumulation of lysosomes and/or hybrid organelles with properties of lysosomes, late endosomes and autophagosomes in response to JIP3 loss-of-function mutations in multiple model organisms (Drerup and Nechiporuk, 2013; Edwards *et al*., 2013; Gowrishankar *et al*., 2017; Hill and Colon-Ramos, 2019).

Our new results define baseline lysosome properties in human iPSC-derived neurons and show that KO of the human *JIP3* gene results in the accumulation of protease-deficient lysosomes in swollen axons and increased Aβ42 production. Furthermore, we found that this axonal lysosome accumulation is enhanced by the additional KO of *JIP4*, revealing that JIP4, previously shown to regulate lysosome movement in non-neuronal cells (Willett *et al*., 2017), has an overlapping function with JIP3 in regulating axonal lysosome transport in neurons. Our results, which parallel and extend recent observations from rodent models, emphasize the power of human iPSC-derived neurons as a valuable tool for investigating human neuronal cell biology and neurodegenerative disease mechanisms. These new observations are also relevant for the development of models to study recently discovered neurodevelopmental defects arising from mutations in the human *JIP3* (also known as *MAPK8IP3*) gene (Iwasawa *et al*., 2019; Platzer *et al*., 2019).

## Materials and Methods

### CRISPR-Cas9 mediated knock out of JIP3

We used an established WTC-11 human iPSC line (provided by Michael Ward, NINDS) that was engineered to harbor a doxycycline-inducible NGN2 transgene expressed from the AAVS1 safe harbor locus in order to support an efficient protocol for their differentiation into neurons (referred to as i^3^Neurons) with properties of layer 2/3 cortical glutamatergic pyramidal cells (Wang *et al*., 2017; Fernandopulle *et al*., 2018). These iPSCs were grown on Matrigel coated dishes in E8 media (Gibco). For CRISPR-Cas9-mediated gene editing, cells were harvested using accutase (Corning) and 1.5 million cells were resuspended in Mirus nucleofector solution and electroporated with 5 ug of px458 plasmid (Addgene plasmid #48138) containing a small guide RNA (see Table S1 for oligonucleotide information) targeted against the *JIP3* (also known as *MAPK8IP3*) gene using an Amaxa 2D nucleofector. Electroporated cells were then plated into one well of a 24-well plate and GFP-positive cells were selected by FACS after 4 days. Cells were once again plated communally post-sorting and serially diluted 4 days later to yield clonal populations for screening. 10 colonies were selected per sgRNA and screened for mutations using PCR amplification of genomic DNA flanking the sgRNA target site followed by sequencing of the amplicons (see Supplemental Table S1 for primer details).

### Generation of JIP3/4 Double knock out cells

The JIP3 KO iPSC line was subjected to a second round of CRISPR-based editing by the method described above to disrupt the *JIP4* gene in these cells (see Table S1 for oligonucleotide information). PCR amplification of genomic DNA flanking the sgRNA target site followed by sequencing of the amplicons (see Supplemental Table S1 for primer details), was carried out to identify cells harboring the double knock out.

### Generation of stable iPSC line expressing LAMP1-GFP

LAMP1-GFP was amplified using PCR (using primers listed in Table S2) from Addgene plasmid #34831 and then cloned into lentiviral vector FUGW (Addgene plasmid #14883) from which the GFP fragment had been excised, by digesting the vector with BamHI and EcoRI. FUGW was a gift from David Baltimore (Addgene plasmid # 14883; http://n2t.net/addgene: 14883; RRID:Addgene_14883). LAMP1-mGFP was a gift from Esteban Dell’Angelica (Addgene plasmid # 34831; http://n2t.net/addgene: 34831; RRID:Addgene_34831).

HEK 293 FT cells were transfected with the newly made LAMP1-GFP lentiviral vector, along with the psPAX2 (Addgene #12260) and PCMV (Addgene #8454) packaging plasmids. pCMV-VSV-G was a gift from Bob Weinberg (Addgene plasmid # 8454; http://n2t.net/addgene:8454; RRID:Addgene_8454). psPAX2 was a gift from Didier Trono (Addgene plasmid # 12260; http://n2t.net/addgene:12260; RRID:Addgene_12260). The supernatant from these cells containing the virus was collected, concentrated, and then diluted into E8 medium containing the Y-27632 ROCK inhibitor (Tocris). This was added to freshly split iPSCs (1.5 million cells in one well of a 6-well plate). The media on the cells was changed after 48 hours. These cells were then expanded and LAMP1-GFP expressing cells were selected by FACS.

### ELISA-based measurements of Aβ42 in i^3^Neurons

I^3^Neurons were differentiated from Control and JIP3 KO iPSCs as described previously (Fernandopulle *et al*., 2018). i^3^Neurons were grown on Poly-Ornithine (PO; Sigma Aldrich) coated 6-well plates for 21 days before Aβ42 ELISA measurements were performed as per a previously validated ELISA assay for the measurement of human Aβ42 (Teich *et al*., 2013). In brief, the cells were lysed into Tris-EDTA and then processed for ELISA based measurements of Aβ42 using a high-sensitivity ELISA kit (Wako; 292–64501). Standard curves were generated using Aβ1-42 peptide standards for each set of experiments.

### Immunofluorescence analysis of i^3^Neurons

I^3^Neurons differentiated for one to six weeks on 35 mm Mattek glass bottom dishes were processed for immunostaining as described previously (Gowrishankar *et al*., 2017). See Table S2 for antibody information.

### Immunoblotting experiments

I^3^Neurons (Control and hJIP3 KO) were grown on PO-coated 6-well plates (500,000 cells/well). After 14 or 21 days of differentiation, i^3^Neurons were washed with ice-cold PBS and then lysed in lysis buffer [Tris-buffered saline (TBS) with 1% Triton, protease inhibitor cocktail and phosphatase inhibitor] and spun at 13,000 *g* for 5 minutes. The supernatant was collected and incubated at 95°C for 5 minutes in SDS sample buffer before SDS-PAGE, transfer to nitrocellulose membranes, and immunoblotting. For immunoblotting experiments comparing iPSCs with i^3^Neurons, lysates of the parental iPSC line (∼70% confluent) were compared to the lysates from i^3^Neurons differentiated from these cells. See Table S2 for antibody information.

### Microscopy

Standard confocal images were acquired using a Zeiss 880 Airyscan confocal microscope via a 100X plan-Apochromatic objective (1.46 NA) with 2X optical zoom. Live imaging of lysosome dynamics in i^3^Neurons was carried out using Airyscan imaging mode using 100X objective with 1.5x or 2x optical zoom and scan speeds of 1-2 frames per second. Zeiss Zen software was used for processing of the Airyscan images. Further image analysis was performed using FIJI/ImageJ software (Schindelin *et al*., 2012).

### Analysis of lysosome properties

To measure the mobility of LAMP1-positive and Lysotracker-positive organelles, i^3^Neurons stably expressing LAMP1-GFP were incubated with 50nM Lysotracker Deep Red for 3 minutes and then rinsed gently twice with warm (37°C) imaging medium (136 mM NaCl, 2.5 mM KCl, 2 mM CaCl_2_, 1.3 mM MgCl_2_, and 10 mM HEPES, pH 7.4, supplemented with BSA and glucose) and imaged at 37°C in the same buffer for 10 minutes in Airyscan mode as described above. For these live imaging studies, neurons were grown at low densities and axons were identified based on their length, tracing the longest neurite. Kymographs were generated from the videos using the “reslice” function on “straightened” axons (ImageJ/FIJI), and the percentage of Lysotracker-positive LAMP1 organelles, per axon, was counted. The dynamics and directionality of LAMP1 organelles with respect to Lysotracker-positive vesicles was measured using the kymographs. Organelles were considered stationary if they moved <2 µm in 2 minutes.

### Image analysis for measurement of lysosome protein enrichment

i^3^Neurons were co-stained for LAMP1 and either Cathepsin B, L or D and imaged by laser-scanning confocal microscopy (100X objective). Z-stacks were acquired to encompass entire soma and neurites. ImageJ/FIJI software was used to calculate the enrichment of the respective lysosomal proteins in the soma versus neurites. To this end, regions of interest in each image were outlined for LAMP1-positive endo-lysosomes in neurites and the cell body. The mean intensity in each such region was determined, and the ratio between them (neurite to soma lysosomes) was calculated for each lysosomal protein. The mean intensity in endo-lysosomes in neurites was expressed as a ratio to the average mean intensity obtained from endo-lysosomes within the corresponding neuronal cell body.

### Statistical analysis

Data are represented as mean ± SEM unless otherwise specified. Statistical analysis was performed using Prism 8 software. Detailed statistical information (specific test performed, number of independent experiments, and p-values) is described in the respective Figure legends.

## Results

To evaluate the suitability of the i^3^Neuron model (Wang *et al*., 2017; Fernandopulle *et al*., 2018); (Figure 1A) for the investigation of neuronal lysosome biology, we began with a characterization of key properties of these organelles based on recent findings from studies of lysosome distribution and maturation in mouse neurons and brain tissue (Gowrishankar *et al*., 2015; Farias *et al*., 2017; Cheng *et al*., 2018; Yap *et al*., 2018). As we had previously observed in mouse neurons both *in vitro* and *in vivo* (Gowrishankar *et al*., 2015), LAMP1-positive organelles in the i^3^Neurons were more abundant in the soma, as compared to dendrites and axons (Figures 1B, S1A) and the LAMP1-positive organelles localized in the soma of i^3^Neurons were more enriched in cathepsins L, D and B (luminal proteases) compared to those localized in axons and dendrites (Figure 1C-E; Figure S1 B, C). We also established that these i^3^Neurons polarize and establish axons and make synapses by staining for an axon initial segment marker, TRIM46 (Figure S1D) and synapsin I, a presynaptic protein (Figure S2A). This relationship between the abundance of luminal hydrolases that correlates with lysosome subcellular positioning in neurons likely reflects a spectrum of maturation between late endosomes, autophagosomes and lysosomes (Gowrishankar *et al*., 2015; Farias *et al*., 2017; Cheng *et al*., 2018; Yap *et al*., 2018). Consistent with lysosomal hydrolases being enriched within lysosomes in the soma, most of the fluorescence from DQ-BSA, an endocytic cargo whose fluorescence is de-quenched following cleavage by lysosomal proteases (Marwaha and Sharma, 2017), was observed in the soma of i^3^Neurons (Figure S1E). For simplicity, and reflecting the eventual degradative function mediated by this pathway, we will subsequently refer to this collection of closely related organelles as lysosomes (Ferguson, 2018).

**Figure 1:**
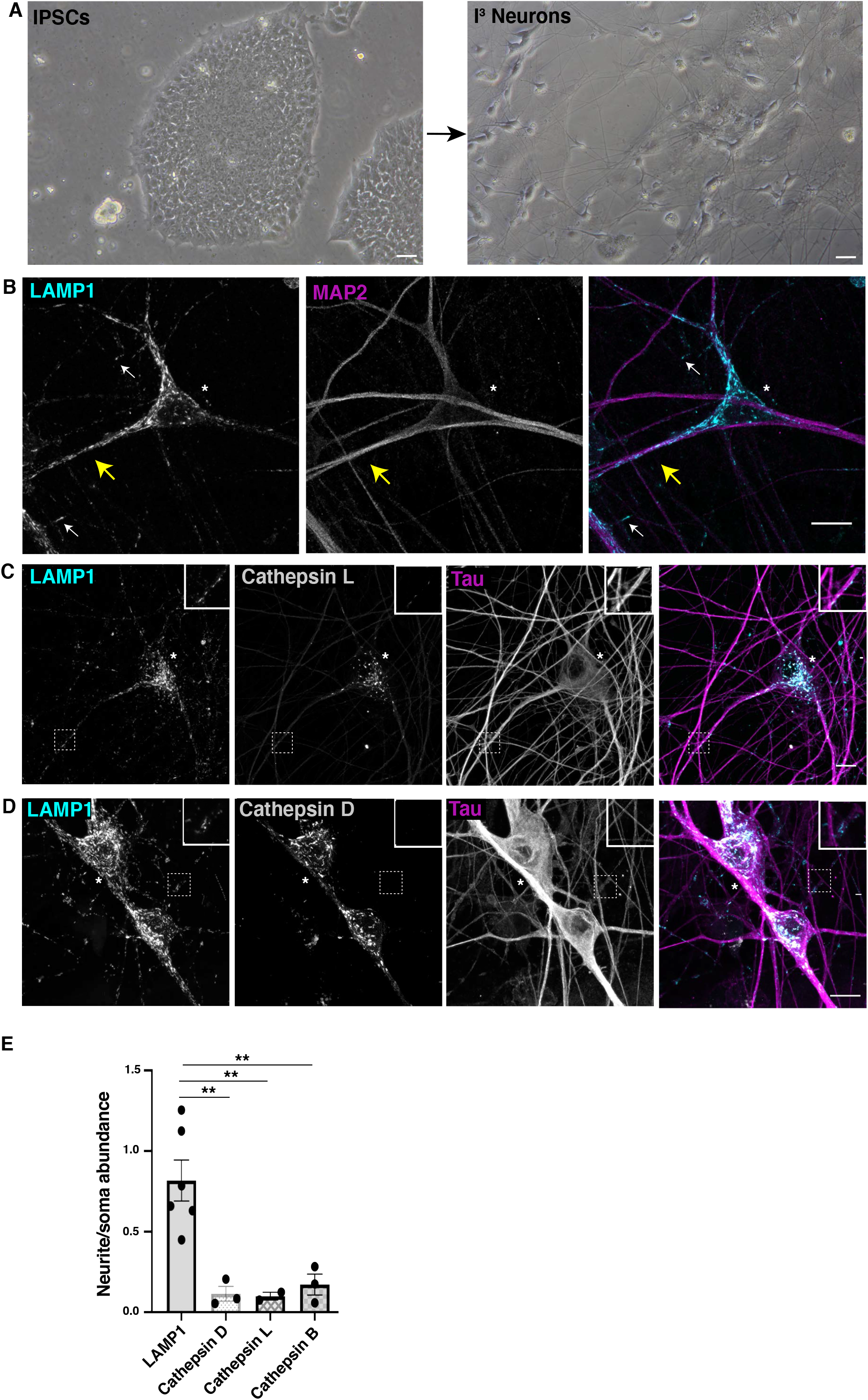
Cathepsin-rich lysosomes are concentrated in the soma of i^3^Neurons. A. Phase contrast images of an iPSC colony (left) and i^3^Neurons differentiated for two weeks (right). Scale Bar, 10μm B. Confocal immunofluorescence image of an i^3^Neuron stained for LAMP1 (cyan) and MAP2 (magenta), showing concentration of LAMP1-positive organelles in the soma (asterisk) when compared to the MAP2-positive neurites (yellow arrows). White arrows point to LAMP1 vesicles in MAP2-negative axons. Scale Bar, 10μm. C. Immunofluorescence staining of i^3^Neurons for cathepsin L (grey), LAMP1 (cyan) and Tau (magenta), showing that LAMP1-postive organelles in neurites are relatively deficient of cathepsin L compared to those in the neuronal cell body. Scale Bar, 10μm. Inset scale bar,1 μm. D. Immunofluorescence staining of i^3^Neurons for cathepsin D (grey), LAMP1 (cyan) and Tau (magenta), showing that LAMP1-postive organelles in neurites are relatively deficient in cathepsin D compared to those in neuronal cell bodies. Scale Bar, 10μm. Inset scale bar, 1μm. E. Quantification of the relative enrichment of lysosomal proteins in neurites compared with neuronal cell body lysosomes (mean ± SEM). At least 2 to 3 different biological replicates (independent differentiation) were analyzed for each condition. \*\**P* < 0.01; (ANOVA with Dunnett’s multiple comparisons test).

In order to visualize lysosomes in live cell imaging studies, we generated iPSCs that stably express GFP-tagged LAMP1 (Eskelinen, 2006). This strategy yielded expression levels that were comparable to the levels of endogenous LAMP1 (Figure 2A). LAMP1-GFP-positive organelles were less abundant in axons when compared to the soma or dendrites (Figure 2B; Figure S2A), similar with what is observed with endogenous LAMP1 staining in the i^3^Neurons (Figure 1B; white and yellow arrows highlight LAMP1 in axons and dendrites respectively). They also exhibited bi-directional movement (Video 1). Since LAMP1 localizes to a spectrum of endo-lysosomal organelles (late endosomes, lysosomes as well as biosynthetic intermediates (Swetha *et al*., 2011; Pols *et al*., 2013; Cheng *et al*., 2018), we next imaged the dynamics of axonal LAMP1-GFP-positive organelles from cells that had additionally been labeled with Lysotracker, a dye that more selectively accumulates with the lumen of lysosomes due to their acidic pH. These experiments revealed distinct populations of LAMP1-positive organelles in axons of Control i^3^Neurons based on differences in their motility and acidification. Firstly, only ∼20% of the LAMP1 organelles in the axon were Lysotracker-positive (Figure 2C D,). Interestingly, these LAMP1 and lysotracker double positive organelles moved predominantly in the retrograde direction (Figure. 2C; Videos 2-4). In contrast, when the directionality of all LAMP1-positive organelles was examined, we observed that a slight majority of LAMP1 vesicles moved in the anterograde direction (Figure. 2E; Video 2-4). These anterogradely-moving organelles that do not label with lysotracker (Videos 2-4) may represent Golgi-derived biosynthetic, LAMP1-containing vesicles that deliver newly made lysosomal proteins for the maturation of endosomes and autophagosomes into lysosomes within the axon (Swetha *et al*., 2011; Pols *et al*., 2013).

**Figure 2:**
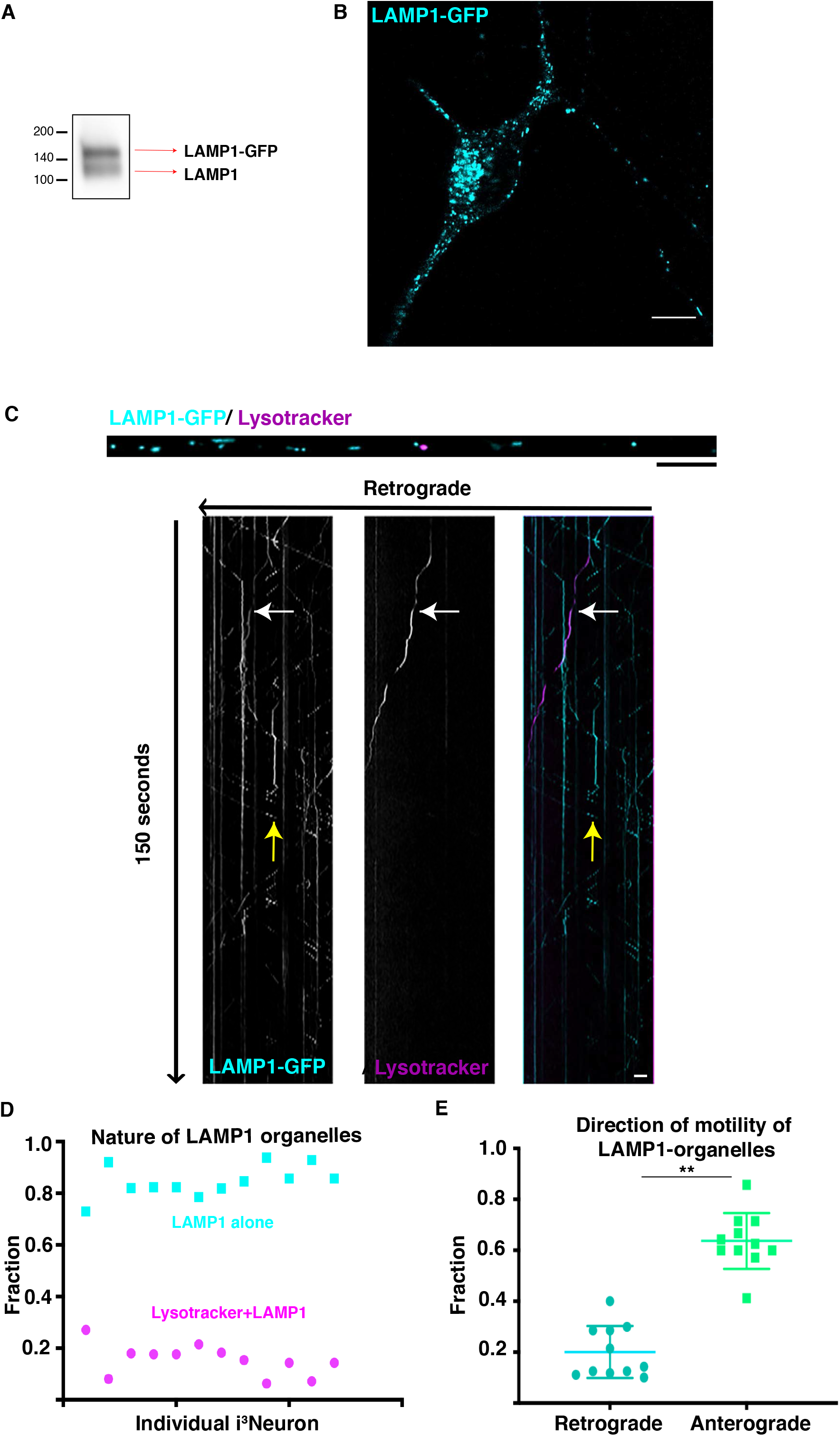
Acidification and movement of LAMP1-positive organelles in i^3^Neurons. A. Immunoblot showing levels of LAMP1-GFP and endogenous LAMP1 in lysates from an i^3^Neuron culture that was differentiated from an iPSC line that stably expresses a LAMP1-GFP transgene. B. LAMP1-GFP subcellular distribution aligns well with endogenous LAMP1 in i^3^Neurons (10 days of differentiation). Scale Bar, 10μm. C. Fluorescence image of a portion of an axon, ‘straightened’ using ImageJ, from an i^3^Neuron expressing LAMP1-GFP (cyan) and stained with Lysotracker-Deep Red (magenta). Scale Bar, 5μm. D. Kymograph of lysotracker (magenta) and LAMP1-GFP (cyan) dynamics in an axon, from a video taken over 2.5 minutes at 1 frame per second. Scale Bar, 5μm. White arrow: Lysotracker-positive+LAMP1 double positive vesicle. Yellow arrow: small, anterogradely moving LAMP1 vesicle. E. Quantification of the fraction of axonal lysotracker and LAMP1-positive organelles in individual neurons. Data reflects 157 LAMP1 organelles analyzed from a total 12 i^3^Neurons analyzed from 3 independent experiments F. Plot summarizing the direction of movement of LAMP1-positive organelles. Each point on the graph represents the fraction of LAMP1 organelles moving in that direction from a single neuron. Data reflects LAMP1 organelles from 11 i^3^Neurons from 3 independent experiments. (p<0.01, unpaired t test).

Having established these baseline properties of i^3^Neuron axonal lysosomes, we next tested how they responded to a JIP3 loss-of-function mutation, a genetic perturbation which promotes axonal lysosome accumulation in other neuronal model systems (Drerup and Nechiporuk, 2013; Edwards *et al*., 2013; Gowrishankar *et al*., 2017). Consistent with past reports of its neuronal enrichment (Gowrishankar *et al*., 2017), the JIP3 protein was undetectable in iPSCs but highly abundant in the i^3^Neurons (Figure 3A). Meanwhile, LAMP1 was present at comparable levels in both iPSCs and i^3^Neurons [although it exhibited a difference in mobility between iPSCs and i^3^Neurons (Figure 3B)]. This mobility shift could reflect differences in the extent of glycosylation of LAMP1 in neuronal cells as compared to non-neuronal cells and might have an impact on the ability of LAMP1 to serve as a transporter for the efflux of cholesterol from neuronal lysosomes (Li and Pfeffer, 2016).

**Figure 3:**
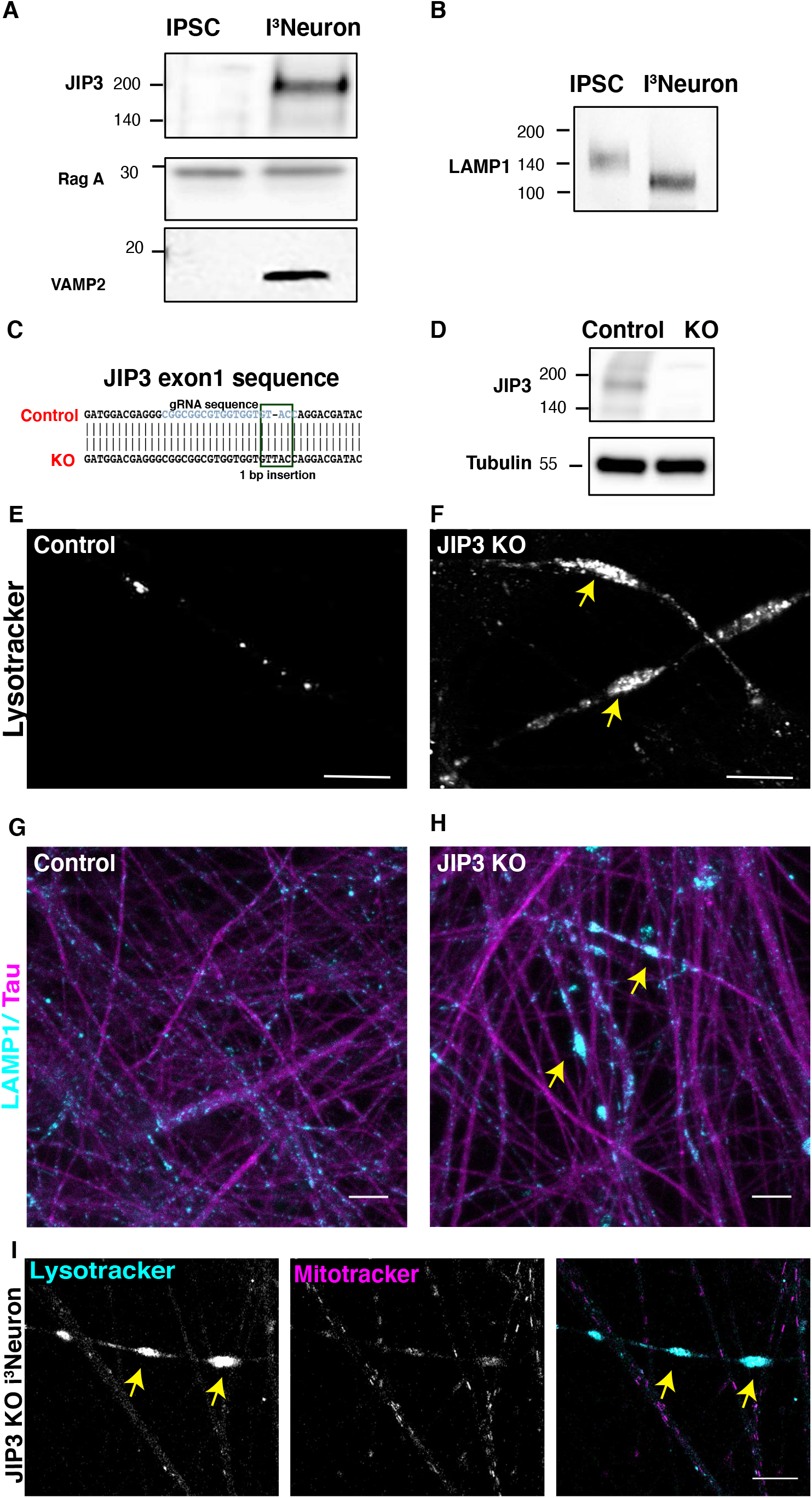
Lysosomes accumulate within axonal swellings in JIP3 KO i^3^Neurons. A. Immunoblots of samples from the parental iPSC line and differentiated i^3^Neurons (21 days of differentiation). B. Immunoblot for LAMP1 in the parental iPSC line and differentiated i^3^Neurons. C. Alignment of a portion of the exon 1 sequence for human JIP3 that shows the location of a single base-pair insertion in the KO cells. The sgRNA target sequence is highlighted in blue. D. JIP3 immunoblot reveals the complete loss of the protein in CRISPR-edited JIP3 KO i^3^Neurons (21 days of differentiation), while it is abundant in the i^3^Neurons differentiated from parental “Control” iPSCs. E and F. Fluorescence images of Lysotracker Red DND-99-stained Control and JIP3 KO i^3^Neurons grown at low density (12 days of differentiation). Yellow arrows highlight axonal swellings filled with lysosomes in the JIP3 KO neurons. G and H. Immunofluorescence image of Control (G) and JIP3 KO (H) i^3^Neurons (12 days of differentiation) stained for LAMP1 (cyan) and Tau (magenta) showing large LAMP1-positive accumulations in the KO culture(yellow arrows) while relatively fewer vesicles seen in neurites of Control cultures. Scale bar, 50 μm. I. Confocal images of JIP3 KO i^3^Neurons (12 days of differentiation) stained with Lysotracker Deep Red (cyan) and Mitotracker Green (magenta). Although some mitochondria are present in the LAMP1-positive axonal swellings, they are not enriched as compared to the LAMP1 signal. Scale bar, 10 μm.

Having established that JIP3 is highly expressed in the i^3^Neuron model, we next used CRISPR-Cas9 mediated gene editing to generate a JIP3 KO iPSC line and obtained cells with a homozygous 1 bp insertion in exon 1 (Figure 3C). This mutation results in a frameshift and premature stop codon that will prevent JIP3 protein translation and is furthermore predicted to trigger nonsense mediated decay of the mutant transcript. Immunoblotting confirmed the loss of JIP3 protein in i^3^Neurons derived from these JIP3 mutant iPSCs (Figure 3D). Consistent with studies of mouse JIP3 KO cortical neuron primary cultures, the human JIP3 KO i^3^Neurons developed lysosome-filled axonal swellings as revealed by lysotracker labeling (Figure 3E, F) and LAMP1 immunofluorescence (Figure 3 G, H). The axonal localization of lysosome accumulation was confirmed by their localization to tau-positive (Figure 3H), MAP2-negative (Figure S3A, B) neurites. These swellings were observed by 10-12 days of differentiation (Figure 3F) and became more penetrant with age (Figure S3C; 21 days of differentiation). In contrast to lysosomes, mitochondria did not prominently accumulate in these swellings (Figure 3I).

Having established that loss of JIP3 results in axonal lysosome-filled swellings resembling those found at Alzheimer’s disease amyloid plaques (Gowrishankar *et al*., 2015), we next interrogated the role of JIP3 on amyloid precursor protein (APP) processing into Aβ peptides in this human neuron model. Our previous studies had demonstrated that accumulation of axonal lysosomes in JIP3 KO mouse neurons was accompanied by increased levels of BACE1 (β-site APP cleaving enzyme) (Gowrishankar *et al*., 2017), the protease that initiates amyloidogenic processing of APP. We found a similar build-up of BACE1 protein in the human JIP3 KO i^3^Neurons (Figure 4 A, B). JIP3 KO i^3^Neurons also exhibited increased proximity of APP and BACE1 [as measured by reconstitution of fluorescence from split Venus fragments fused to APP and BACE1 respectively (Das *et al*., 2016)] in the axonal swellings in (Figure 4C). In contrast, Control i^3^Neurons had very little punctate signal arising from APP and BACE1 proximity (Figure 4C). This was accompanied by a substantial increase in levels of intracellular Aβ42 peptide in the JIP3 KO cultures (Figure 4D). Axonal lysosome accumulations in the JIP3 KO i^3^Neurons are thus accompanied by increased APP processing.

**Figure 4:**
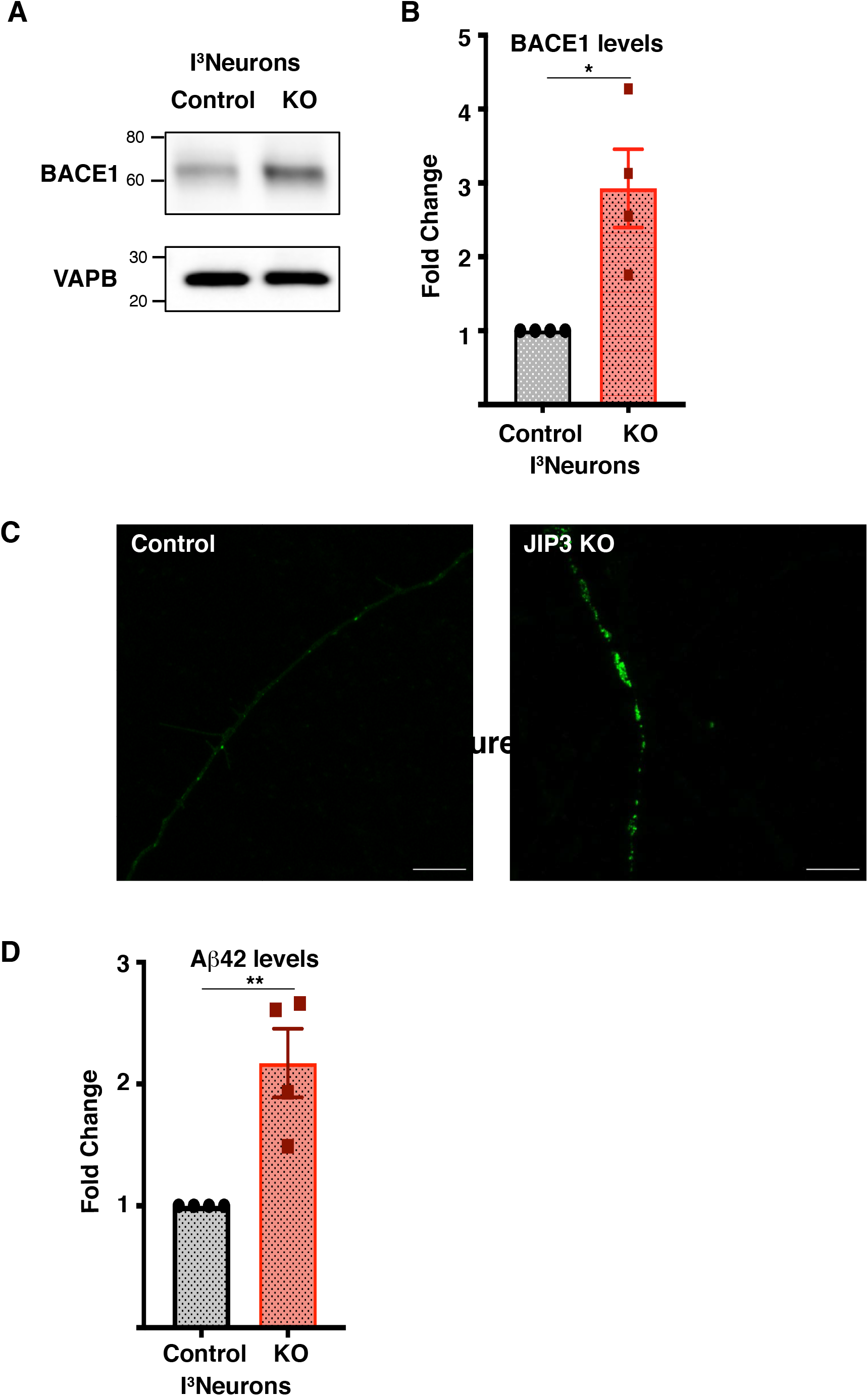
JIP3 KO i^3^Neurons exhibit increased BACE1 and Aβ42 levels. A. Immunoblot reveals increased levels of BACE1 protein in JIP3 KO i^3^Neurons compared to Control (VAPB-loading control) B. Quantification of BACE1 levels in JIP3 KO i^3^Neurons compared to Control (n=4; mean ± SEM; p<0.05; unpaired t-test). C. Airyscan confocal images of Control and JIP3 KO axons expressing APP and BACE proteins tagged with split Venus fragments (APP: VN+BACE-1: VC) showing accumulation of the signal arising from their proximity within axonal swellings in JIP3 KO i^3^Neurons. D. Quantification of endogenous human Aβ42 levels based on ELISA measurements from n=4 independent Control and JIP3 KO i^3^Neuron cultures differentiated for 21 days (mean ± SEM; p<0.01, unpaired t test).

Although the JIP3 KO phenotypes are striking, the impact on axonal lysosome accumulation is age dependent (Figure S3C). This indicates the existence of redundancy in mechanisms for lysosome axonal transport such that JIP3 only becomes critical as the neurons mature. Human JIP4 was previously proposed to play a role in the dynein-dependent movement of lysosomes from the periphery to perinuclear region, in non-neuronal cells (Willett *et al*., 2017). Studies in cultured neurons from JIP3+JIP4 double KO mice indicated a functional redundancy in these molecules for kinesin-dependent axonal transport (Sato *et al*., 2015). However, the impact on lysosomes was not investigated in these double KO neurons. Due to sequence conservation between JIP3 and JIP4 and the ability of JIP4 to promote lysosome movement in other cell types, we next examined the effect of knocking out both JIP3 and JIP4 in our human iPSC model system. Cas9-based editing of JIP4 in the JIP3 mutant iPSC line resulted in a 4 bp deletion in exon 9 of JIP4 (Figure 5A). iPSCs lacking both JIP3 and JIP4 were viable and could still be differentiated into i^3^Neurons. Immunoblotting confirmed the loss of both proteins in i^3^Neurons differentiated from this JIP3+4 double knock out (DKO) iPSC line (Figure 5B). Confocal microscopy revealed that the JIP3+JIP4 double KO cells exhibited axonal lysosome accumulations that were far more severe than those observed in the JIP3 single KO (Figure 5 C-I).

**Figure 5:**
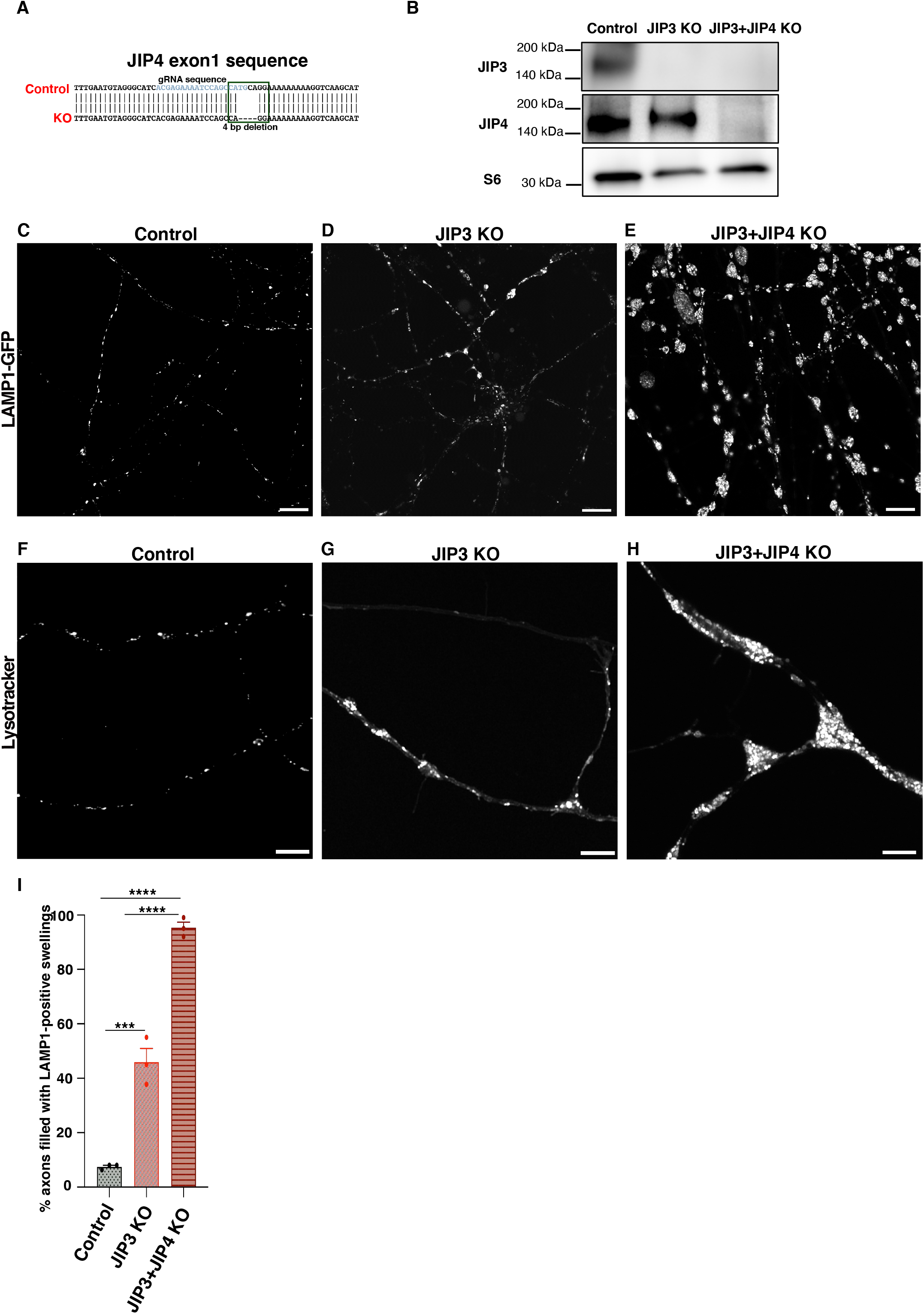
JIP3/4 DKO i^3^Neurons exhibit exacerbation of lysosomal phenotype compared to JIP3 single KO neurons. A. Alignment of a portion of the exon 9 sequence for human JIP4 that shows the location of a 4 base-pair deletion in the DKO cells. The sgRNA target sequence is highlighted in blue. (B) Immunoblot showing levels of JIP3 and JIP4 in Control, JIP3 KO and JIP3+JIP4 DKO i^3^Neurons differentiated for 13 days; ribosomal protein S6 (S6) was used as a loading control. C-E. Confocal image of Control i^3^Neurons, JIP3 KO i^3^Neurons and JIP3+JIP4 i^3^Neurons (DIV 13), stably expressing LAMP1-GFP. Scale bar, 10 μm. Insets highlight lysosome accumulations in the KO neurons. F-H. Confocal image of Control i^3^Neurons,JIP3 KO i^3^Neurons and JIP3+JIP4 DKO i^3^Neurons (DIV 13) stained for Lysotracker. I. Quantification of % axons of Control i^3^Neurons, JIP3 KO i^3^Neurons and JIP3+JIP4 i^3^Neurons showing LAMP1-filled axonal swellings. \*\*\**P* < 0.001; \*\*\*\**P* < 0.0001 (ANOVA with Dunnett’s multiple comparisons test).

## Discussion

In this study, we used human i^3^Neurons as a model for cell biological and genetic studies of axonal lysosome traffic with potential human disease relevance. Our results from JIP3 KO i^3^Neurons reveal potential Alzheimer’s disease relevant consequences of lysosome axonal transport defects. We further generated JIP3+JIP4 double KO lines and established that JIP3 and JIP4 proteins have over-lapping functions in promoting the axonal transport of lysosomes. In addition to these discoveries, our data strengthens confidence that this human iPSC-derived neuronal model can serve as a valuable alternative to traditional rodent neuron primary culture models for the investigation of neuronal cell biology, including the transport of endo-lysosomal and autophagic organelles, as also suggested by other recent studies (Liao *et al*., 2019; Boecker *et al*., 2020).

Our experiments additionally yielded several insights relating to heterogeneity of the axonal LAMP1-positive organelles. First, only a small fraction of them (∼20%) were labeled with the lysotracker dye (indicative of an acidic lumen). Second, these lysotracker-positive LAMP1 vesicles were less mobile than the overall pool of LAMP1-positive organelles and their movement was most frequently in the retrograde direction (Videos 2-4; Figure 2C-E). Third, non-acidified LAMP1 organelles were biased toward movement in the anterograde direction and were generally smaller in appearance. Collectively, these observations are consistent with a model wherein the majority of the anterograde LAMP1-positive organelles represent precursors, which deliver LAMP1 and other cargos into the axon. Distinct identities for anterograde versus retrograde axonal LAMP1-positive organelles is consistent with electron microscopy analysis of axonal organelle morphology proximal and distal to a localized blockade of transport where organelles with late endosome/lysosome morphology preferentially accumulated distal to the block which reflects a predominantly retrograde movement (Tsukita and Ishikawa, 1980). More recently, LAMP1-positive anterogradely moving organelles that do not have degradative properties, but which also deliver synaptic vesicle proteins, were reported (Vukoja *et al*., 2018). Our results define LAMP1 vesicle sub-populations that can be discriminated based on their apparent size, movement direction and lysotracker labeling affinity. These observations add to previous studies, which documented a gradient of lysosome acidification along the axon with a higher frequency of acidified lysosomes closer to the cell body (Overly *et al*., 1995; Overly and Hollenbeck, 1996; Maday *et al*., 2012). However, differences in probe pH sensitivity could explain why other investigators using pHluorin (pKa∼7) fused to LAMP1 concluded that essentially all LAMP1 organelles are acidified (Farias *et al*., 2017). The growing appreciation for heterogeneity in axonal endo-lysosome sub-populations raises a host of questions about regulatory mechanisms and physiological functions (Farias *et al*., 2017; Cheng *et al*., 2018; Ferguson, 2018). The experimental tractability of the i^3^Neuron systems for imaging and genetic manipulations makes it an ideal tool for answering these questions.

The increased abundance of BACE1 and Aβ42 peptide that accompanies lysosome transport defects in the JIP3 KO i^3^Neurons parallels past observations in mouse JIP3 KO neuronal primary cultures, transgenic Alzheimer’s disease mouse models as well as human Alzheimer’s disease brains (Fukumoto *et al*., 2002; Holsinger *et al*., 2002; Zhao *et al*., 2007; Zhang *et al*., 2009; Gowrishankar *et al*., 2017). These observations establish the utility of the i^3^Neurons model for investigating the mechanistic basis of the relationships between lysosome axonal transport and APP cleavage as well as for the development of new strategies to modulate these processes for therapeutic purposes. In addition to its relevance to Alzheimer’s disease, this model will be valuable for understanding mechanisms underlying certain sub-types of hereditary spastic paraplegia that have changes in axonal lysosome abundance associated with them (Edmison *et al*., 2021).

In addition to serving as a model for investigating Alzheimer’s disease, the JIP3 KO i^3^Neuron model may support the investigation of other human neurological diseases. Of most obvious relevance, *de novo* heterozygous loss-of-function variants in the human *JIP3/MAPK8IP3* gene were recently identified as a cause of neurodevelopmental defects resulting in intellectual disability (Iwasawa *et al*., 2019; Platzer *et al*., 2019). The emergence of a severe human neurodevelopmental disease from what is predicted to be just a partial reduction in JIP3 expression suggests an exquisite sensitivity to gene dosage at critical developmental stages. Consistent with this interpretation, large scale exome sequencing revealed strong selective pressure against predicted deleterious variants in the human *MAPK8IP3* gene (Lek *et al*., 2016). Thus, in addition to our original focus on the role of JIP3 in lysosome transport and APP processing mechanisms, there are now broader questions about how JIP3 contributes to human neuronal development that can potentially be answered in the i^3^Neuron model system.

## Acknowledgements

This research was supported in part by the Dementia Discovery Foundation (SMF and PDC), the NIH (NS36251 to PDC; AG062210 TO SMF), and the Kavli Foundation (PDC). S.G. was a recipient of a BrightFocus Foundation postdoctoral fellowship. Competing Interests: PDC is on the Scientific Advisory Board of Casma Therapeutics.

