## Supplemental Figures and legend for "Overlapping roles of JIP3 and JIP4 in promoting axonal transport of lysosomes in human iPSC-derived neurons"

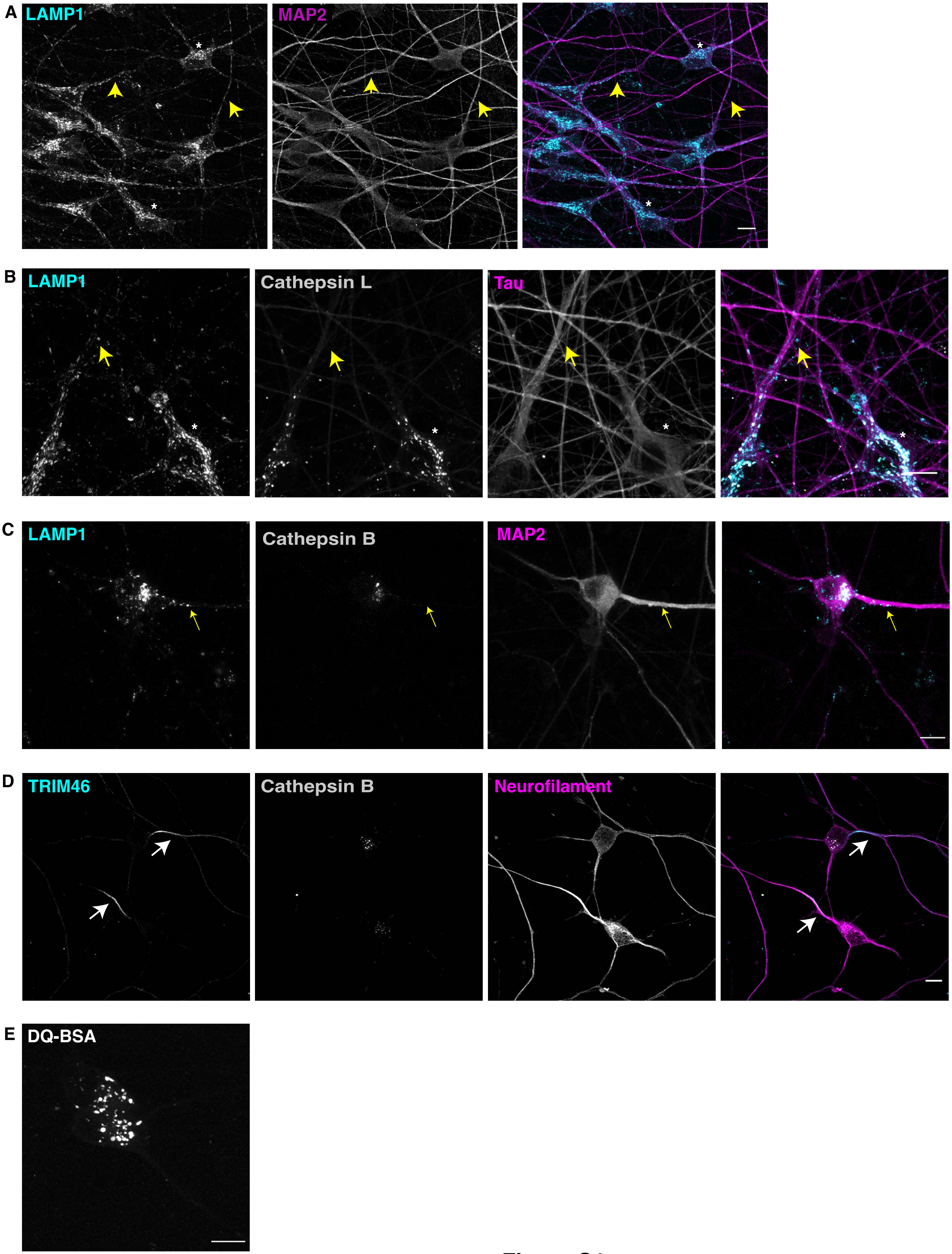
A
LAMP1
MAP2
B
LAMP1
Cathepsin L
Tau
C
LAMP1
Cathepsin B
MAP2
D
TRIM46
Cathepsin B
Neurofilament
E
DQ-BSA

**Figure S1**

**Figure S1: Subcellular distribution of lysosomes in i$^3$Neurons.**

A. Stitched confocal image of i$^3$Neurons (3 weeks of differentiation) co-stained for LAMP1 (cyan; endo-lysosomal compartments) and MAP2 (magenta; dendrites). Scale Bar, 10μm. B. Confocal image of immunofluorescence staining of i$^3$Neurons for cathepsin L (grey), LAMP1 (cyan) and Tau (magenta), showing that LAMP1-postive organelles in neurites (yellow arrow) are relatively deficient of cathepsin L compared to those in the neuronal cell body. C. Confocal image of immunofluorescence staining of i$^3$Neurons for cathepsin B (grey), LAMP1 (cyan) and Tau (magenta), showing that LAMP1-postive organelles in neurites (yellow arrow) are relatively deficient of cathepsin B compared to those in the neuronal cell body. D. Confocal image of immunofluorescence staining of i$^3$Neurons differentiated for two weeks for cathepsin B (grey), TRIM46 (cyan) and Neurofilament (magenta), showing that axon-specification (enrichment of TRIM 46 at axon initial segment; white arrows). E. Airyscan image of i$^3$Neurons showing enrichment of DQ BSA fluorescence (read-out of lysosomal degradation) in soma.

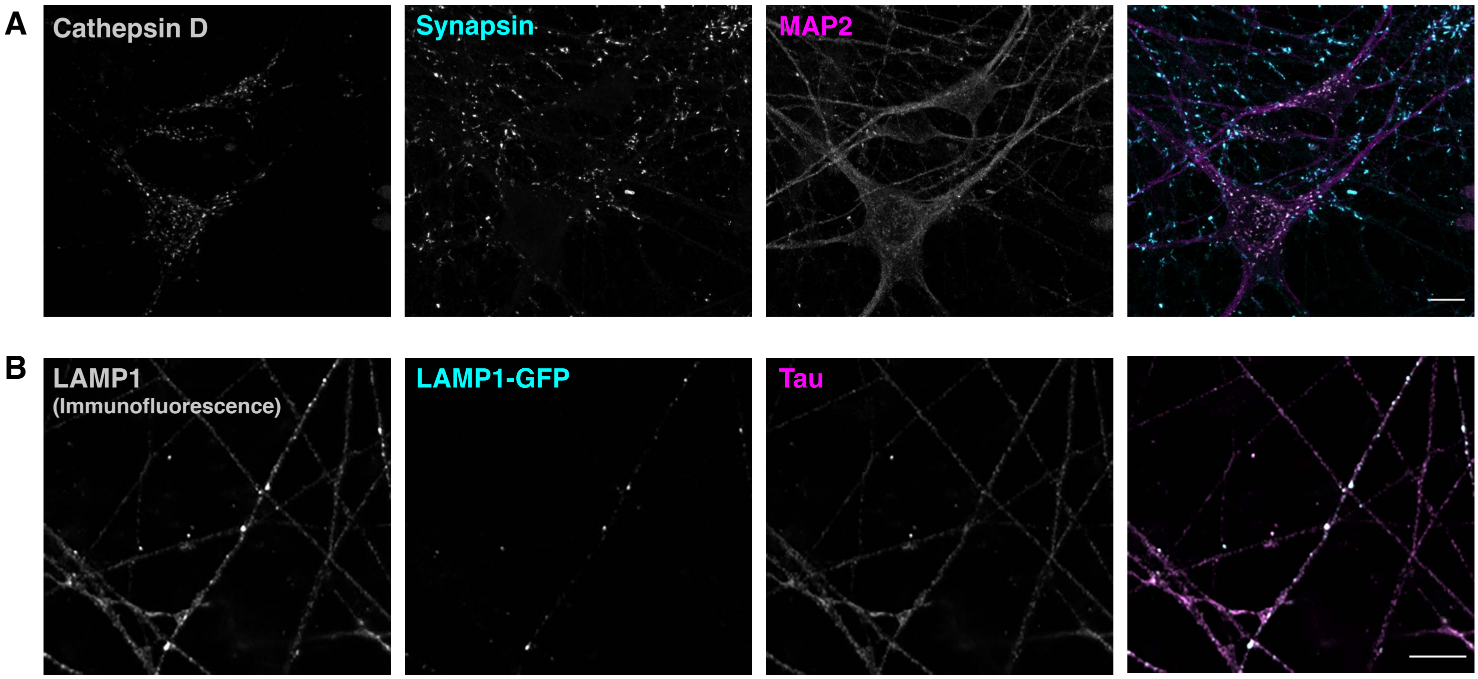
A
Cathepsin D
Synapsin
MAP2
B
LAMP1
(Immunofluorescence)
LAMP1-GFP
Tau

**Figure S2**

**Figure S2: Synapses and LAMP1-GFP in i$^3$Neurons.**

A. Confocal image of immunofluorescence staining of i$^3$Neurons for cathepsin D (grey), Synapsin (cyan) and MAP2 (magenta), showing that synapses form in i$^3$Neurons. B Confocal image of immunofluorescence staining of i$^3$Neurons stably expressing LAMP1-GFP (cyan) for LAMP1(grey), and Tau (magenta), showing the similar distribution of stably expressed LAMP1-GFP to LAMP1.

.

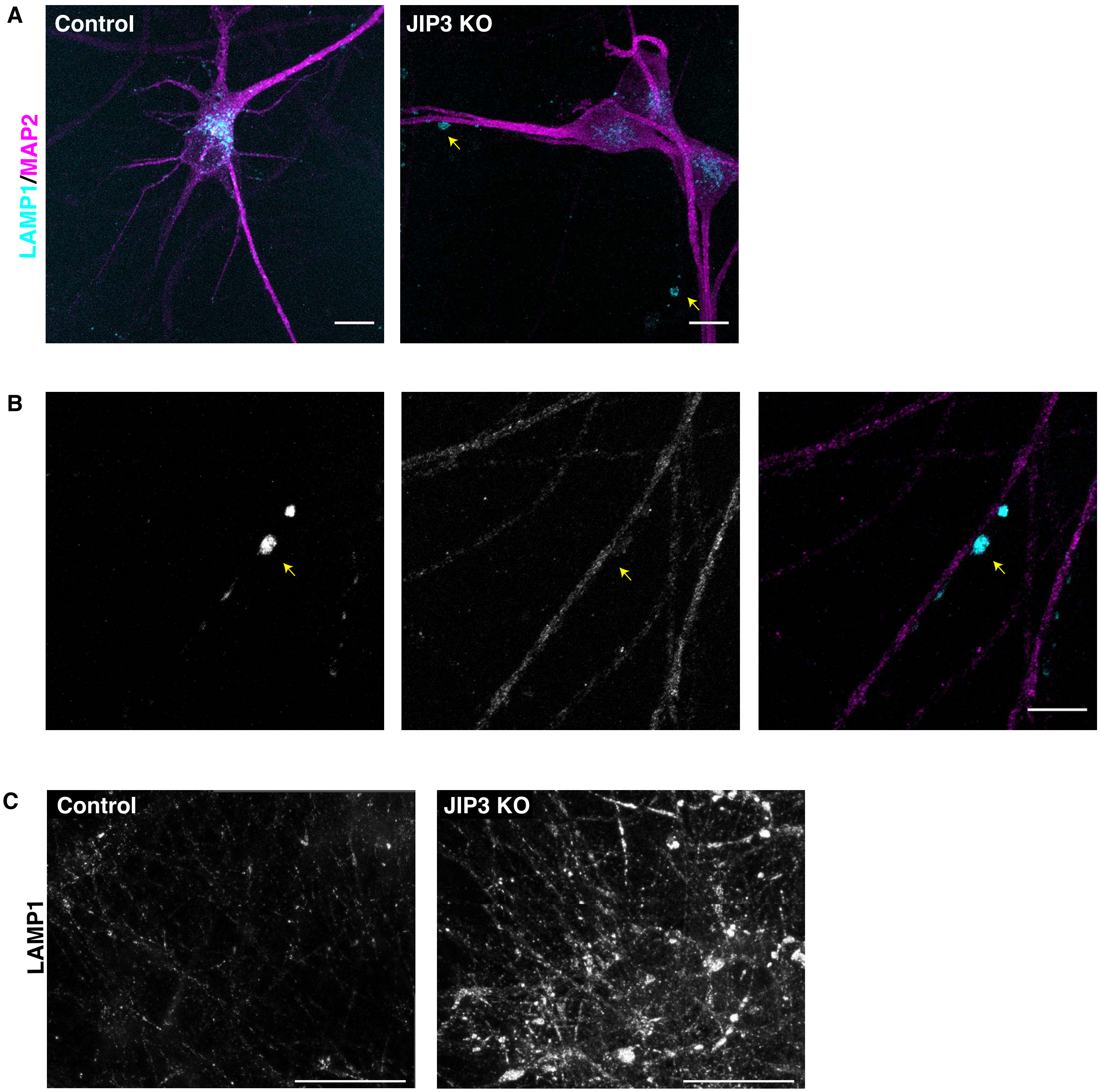


**Figure S3**

**Figure S3: Axonal lysosome accumulations in JIP3 KO i$^3$Neuron.**

A. Immunofluorescence image of Control and JIP3 KO i$^3$Neurons (14 days of differentiation) stably expressing LAMP1-GFP stained for MAP2 showing that lysosomes accumulate in axons (MAP2-negative neurites) in JIP3 KO i$^3$Neurons (yellow arrows). B. Zoomed in image showing LAMP1-vesicle accumulation (cyan) in MAP2-negative axonal swellings in JIP3 KO i$^3$Neurons (yellow arrow). C. Stitched immunofluorescence image of Control and JIP3 KO i$^3$Neurons (21 days of differentiation) showing large LAMP1-positive accumulations in the KO culture. Scale bar, 50 μm.

**Supplemental Video Legends**

**LAMP1 and Lysotracker dynamics in $i^3$Neurons**

Video 1: LAMP1-GFP-positive organelles move bi-directionally in $i^3$Neuron axons. Images were acquired at 2 frames/s, playback rate 21 frames/s. Scale bar, 10 μm.

Videos 2-4: Lysotracker-positive (acidic; Video 3,4) and Lysotracker-negative LAMP1 vesicles (Video 2,4) in axon of an $i^3$Neuron stably expressing LAMP1-GFP and differentiated for a week. Images were acquired at 1 frame/s, playback rate 7 frames/s. Scale bar, 5 μm.

**Table S1: Summary of oligonucleotide primers used in the study**

| **Gene Target** | **Use** | **Sense sequence (5' 3')** | **Antisense sequence** |
|---|---|---|---|
| JIP3 | CRISPR KO | CACCGCGGCGGCGTGGTGGTGTACC | AAACGGTACACCACCACGCCGCCGC |
| JIP4 | CRISPR KO | CCGACGAGAAAATCCAGCCATGC | AAACGTGAGAAAGATGTGCTGCAA |
| JIP3 | Genomic sequencing | ACGCTCGTACTGGGTGA | GCGATGATGGAGATCCAGATG |
| LAMP1 | PCR for cloning of LAMP1-GFP into FUGW lentiviral vector | GGCTGCAGGTCGACTCTAGAGGATCCATGGCGGCCCCC GGCAGCGC | TTATCGATAAGCTTGATATCGAATTCTT ACTTGTACAGCTCGTCCA |

**Table S2: Antibody Summary**

| Antibody | Source | Catalog Number |
|---|---|---|
| BACE1 | Cell Signaling Technology | 5606 |
| Cathepsin B | R&D Systems | AF953 |
| Cathepsin D | R&D Systems | AF1014 |
| Cathepsin L | R&D Systems | AF 952 |
| JIP3 | Sigma | HPA069311 |
| JIP4 | Cell Signaling Technology | 5519 |
| LAMP1 | University of Iowa DSHB | H4A3 |
| LAMP1 | Cell Signaling Technology | 9091 |
| MAP2B | Millipore | AB5622 |
| MAP2 | BD Biosciences | 610410 |
| Neurofilament | Biolegend | SM312-R |
| Rag A | Cell Signaling Technology | 4357 |
| S6 Ribosomal protein | Cell Signaling Technology | 2217 |
| Synapsin 1 | De Camilli Lab | G246 |
| Tau | Cell Signaling Technology | 4019S |
| TRIM46 | Synaptic Systems | 377 003 |
| Tubulin | Sigma | T5168 |
| VAMP2 | Synaptic systems | 104211 |
| VAPB | Sigma Aldrich | HPA013144 |
